# Principles of Computer Numerical Controlled Machining Applied to Cranial Microsurgery

**DOI:** 10.1101/280461

**Authors:** Leila Ghanbari, Mathew Rynes, Jay Jia Hu, Daniel Sousa Shulman, Gregory Johnson, Micheal Laroque, Gabriella Shull, Suhasa B. Kodandaramaiah

## Abstract

Over the last decade, a plethora of tools have been developed for neuroscientists to interface with the brain. Implementing these tools requires precise removal of sections of the skull to access the brain. These delicate cranial microsurgical procedures need to be performed on sub-millimeter thick bone without damaging the underlying tissue and therefore, require significant training. Automating some of these procedures would not only enable more precise microsurgical operations, but also democratize use of advanced neurotechnologies. Here, we describe the ‘Craniobot’, a cranial microsurgery platform that combines automated skull surface profiling with a computer numerical controlled (CNC) milling machine to perform a variety of cranial microsurgical procedures in mice. The Craniobot utilizes a low force contact sensor to profile the skull surface and uses this information to perform micrometer-scale precise milling operations within minutes. We have used the Craniobot to drill pilot holes to anchor cranial implants, perform skull thinning, and open small to large craniotomies. The Craniobot is built using off-the-shelf components for under $1000 and is controlled using open-source CNC programming software.

## INTRODUCTION

The palette of tools available for systems neuroscientists to measure and manipulate the brain has exploded in the last decade. Many *in vivo* neuroscience experiments used simple electrical sensing and stimulation tools, such as tungsten electrodes (Hubel, 1957) and have now evolved to using high channel count silicon-based 3-dimensional (3D) electrodes (Jun et al., 2017; Scholvin et al., 2016). Recent advances in fabrication techniques and materials have enabled the development of flexible neural interfaces (Liu et al., 2015; Viventi et al., 2011; Yazicioglu et al., 2014), injectable neural mesh electrodes (Liu et al., 2015), and 3D organic electrodes (Khodagholy et al., 2016). In parallel, the emergence of optical tools such as optogenetic molecules (Boyden et al., 2005; Chow et al., 2010; Chuong et al., 2014; Zhang et al., 2007), and fluorescent activity reporters (Chen et al., 2013; Dana et al., 2016) have enabled researchers to investigate wide regions of the brain at cellular or near-cellular resolution (Kim et al., 2016; Zong et al., 2017).

As neurotechnologies have advanced, the corresponding cranial microsurgery procedures to deploy them have become more complex. For example, multi-shank neural probes require precisely arrayed craniotomies for insertion into regions of interest. Calcium imaging was initially performed through chronically implanted planar glass coverslips over 3-4 mm diameter craniotomies (Holtmaat et al., 2009). With the emergence of ultra-wide field view imaging technologies, neuroscientists have progressed to implanting coverslips across a whole hemisphere of the cortex (Sofroniew et al., 2016; Stirman et al., 2016) and more recently, across the whole dorsal cortex (Kim et al., 2016). Intact skull preparations are also commonly used for optical imaging on mice, and are particularly useful in studies where neuro-inflammatory effect need to be minimized (Drew et al., 2010; Shih, Mateo, Drew, Tsai, & Kleinfeld, 2012; Silasi, Xiao, Vanni, Chen, & Murphy, 2016). Such procedures require the skull to be thinned down to tens of micrometers.

Mice, the most widely used mammalian model organism, have very thin skulls, typically ranging from 100-700 μm above the dorsal cortex. Any surgical procedure involving removal of bone must be performed with great care and precision, ensuring the underlying meninges and brain tissue are not damaged. Further, the quality of the procedure has a significant bearing on the success of experiments. For instance, *in vivo* patch clamping experiments rely on pristine craniotomy preparations, for high success rate (Desai et al., 2015; Kodandaramaiah et al., 2016, 2018; Margrie et al., 2003). Cranial microsurgery procedures are typically performed using manual tools adapted from dentistry, which are imprecise and require several months of training for a new scientist to become proficient. Consequently, many of the more advanced neurotechnologies end up being confined to a select few lab groups. Automating some of these procedures would potentially enable more precise cranial microsurgeries, facilitate the wider use of advanced neurotechnologies, and potentially open up new kinds of scientific experiments.

Several attempts to automate craniotomies have been made. These include using force or impedance feedback to control the drilling depth (Loschak et al., 2012; Pak et al., 2015; Pohl et al., 2011). Force feedback based systems are typically designed for large animal models and have not been used for automating microsurgical procedures in mice. Impedance sensing feedback (Pak et al., 2015) achieves micrometer scale precision and has been demonstrated in mice, but its performance is affected by the presence of vasculature within the skull and is not generalizable across many areas of the skull. Further, these methods are constrained to complete removal of bone for craniotomies and cannot be used for partial bone removal as in the case of intact skull preparations (Drew et al., 2010; Shih et al., 2012). Other versions include using femtosecond lasers to remove skull tissue (Jeong et al., 2013), which would be impractical to implement in regular neuroscience laboratories due to high cost.

A generalized methodology for performing a wide range of microsurgical procedures is currently not available. If such a methodology can be derived and implemented in an inexpensive robotic system, it could be a valuable resource for systems neuroscientists. Computer Numerical Control (CNC) practices have been ubiquitously used in precision manufacturing and could potentially be adapted to perform these cranial surgical procedures.

Here, we introduce the ‘Craniobot’, a comprehensive microsurgery platform for mice based on a desktop CNC mill. The Craniobot combines automated skull surface profiling with a CNC milling to perform precise microsurgical procedures such as intact skull thinning, small to large craniotomies of arbitrary shapes on the dorsal skull, and drilling pilot holes for anchoring cranial implants. The Craniobot is open-source and is assembled with off-the-shelf components for under $1000.

## RESULTS

### Principles of CNC microsurgery

We hypothesized that if an apparatus could be built to precisely determine the coordinates of pilot points along a cutting path on the skull surface, it would be possible to interpolate a 3D cutting trajectory that can be executed by a CNC mill (**Fig. 1a**). The average thickness of the skull is relatively well conserved within each mouse strain (e.g., C57BL/6J) at a certain age and sex (**Supplementary Fig.1**). Thus, a topographic map of the top surface of the skull is sufficient to perform skull excisions by iteratively milling. The skull can be thinned down to a point where it can be excised without damaging the underlying brain tissue as long as the final depth of milling is less than the minimum thickness of the skull at any point along the desired cutting path. To test this hypothesis, a simple motorized manipulator guided end mill was built and incorporated into a standard rodent stereotax (**Supplementary Fig. 2**). This early prototype allowed us to automate bone tissue milling. The skull surface was profiled manually by lowering the tip of the cutting tool to the skull surface at pilot points along a pre-defined trajectory. Once the contact was confirmed visually through a stereomicroscope, z-coordinate at that point was registered. These registered coordinates were used to interpolate a 3D cutting path for milling the skull. The depth of milling was then iteratively increased until the skull was fragile enough to be excised (**Supplementary Note 1)**. Following the craniotomy, the thickness of the excised skull segments was measured at various locations (**Supplementary Fig. 3)**. The final drilling depth was on average 56.1 ± 30.4 μm (n=13, C57BL/6J mice) and 146.6 ± 25.2 μm (n=6, Thy1-GCaMP6f mice) less than the minimum thickness of the excised skull segment. This confirmed that the iterative milling procedure could successfully perform craniotomies across the skull above the whole dorsal cortex without damaging the underlying tissue.

**Figure 1.**
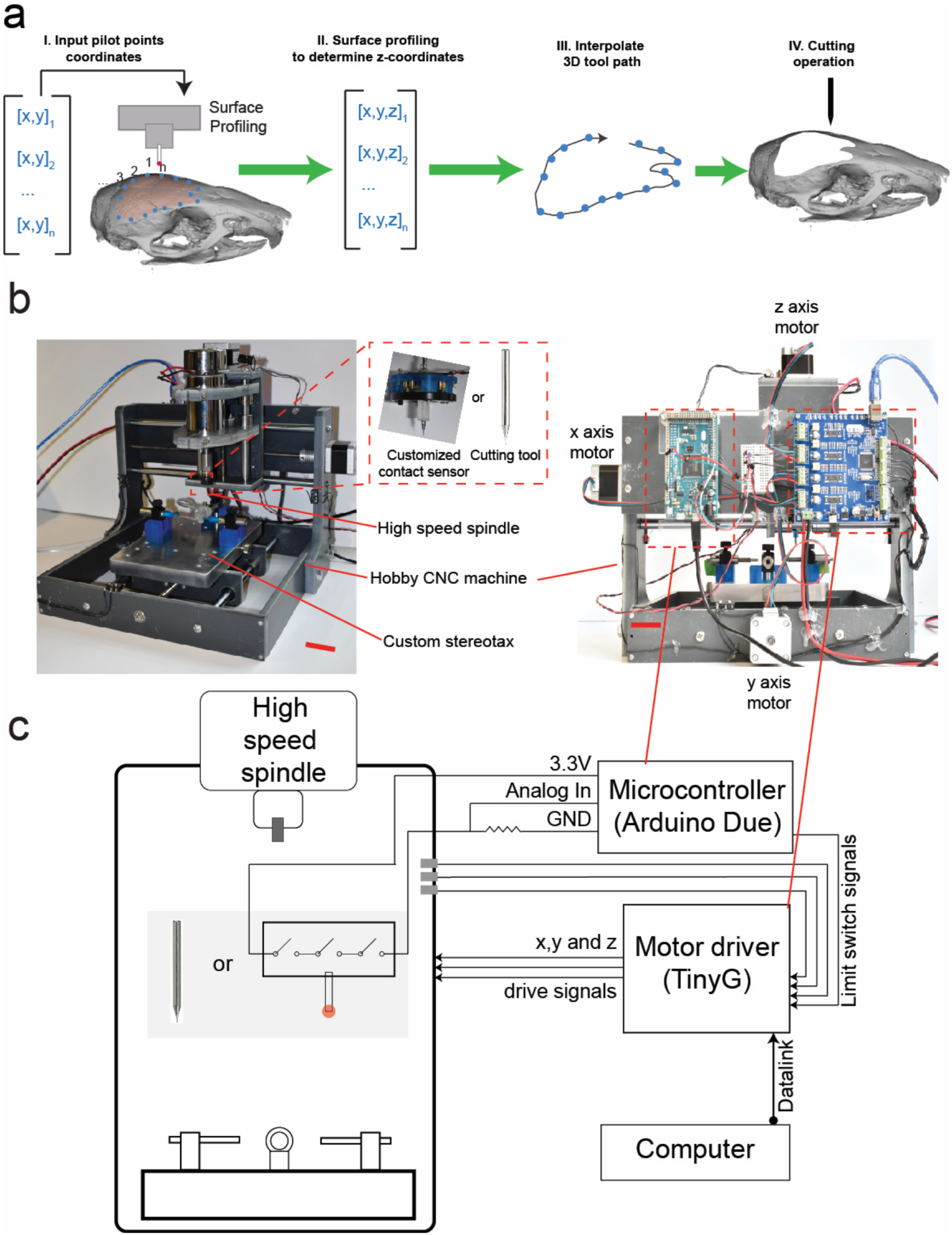
Principle of surface profile guided computer numerically controlled (CNC) skull machining: **(a)** Given a set of x-y coordinates of the pilot points along a desired craniotomy, a device can be used to determine the z coordinates at each point and interpolate a 3D milling path used to perform iterative skull milling. Given a set of x-y coordinates of the pilot points along a desired craniotomy, a device can be used to determine the z coordinates at each point and interpolate a 3D milling path. **(b)** We built the ‘Craniobot’ to perform this algorithm by adapting a CNC milling machine typically used for wood working, to perform automated microsurgical procedures. A 3-axis milling base guides a high-speed spindle which can accommodate either a contact sensor for profiling the skull surface or a cutting tool for machining. It incorporates a custom built stereotax into the mill base. Open-source motor drivers and microcontrollers are integrated to execute the Craniobot algorithm. Scale bar, 4cm. **(c)** A schematic illustrating the electronic components in the Craniobot.

### Craniobot Hardware

We then implemented this milling algorithm along with automated surface profiling in the Craniobot (**Fig. 1b)**. The Craniobot consists of a 3-axis hobby CNC milling machine, typically used by woodworking hobbyists. The milling machine includes a spindle that can either be fitted with a contact sensor or a milling tool. A custom built stereotax was incorporated into the bed of the CNC mill to secure the mouse in the Craniobot (**Supplementary Fig. 4**).

### Automated profiling of skull surface

CNC machining operations rely on precise measurement of substrate surfaces, typically accomplished using a contact sensor with a digital readout. A commercially available contact sensor typically used for profiling metal substrates was modified by incorporating a custom designed linear spring (**Fig. 2a** and **b,** details in **Supplementary Note 2**) to have a 5-10 gf actuation force. This range of actuation force allowed accurate contact detection with the skull surface without any perceptible deformation, as visualized using a stereomicroscope (**Supplementary Video 1**).

**Figure 2.**
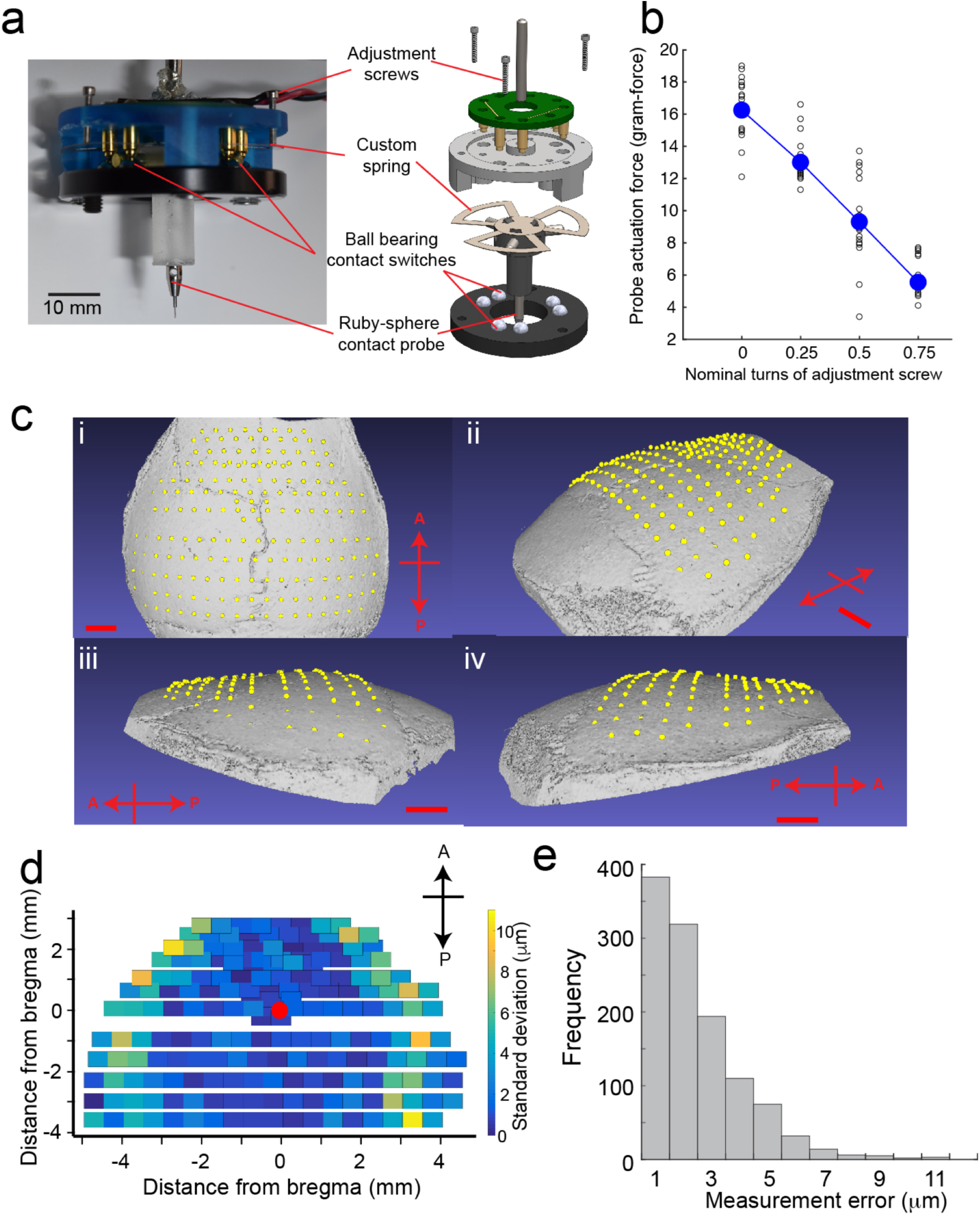
Automatic skull surface profiling (a): We engineered a low force contact sensor for profiling the surface of the skull. Incorporating a custom spring (**Supplementary Note 2**) enabled the contact sensor to accurately detect the skull surface without causing any visible deflection when observed through a stereomicroscope at 60X (**Supplementary Movie 1**) **(b)** Relationship between spring actuation force required to detect contact with the skull and the nominal turns of the adjustment screws. The actuation force ranged from 5 to 10 gf in all experiments. The average actuation force from 20 measurements is shown for each quarter turn of the adjustment screws as a filled circle along with the individual measurements as hollow circles. **(c)** The average point-cloud generated by surface profiling the dorsal skull of an 8-week old male C57BL/6J mouse with a needle-tipped stylus at 192 points overlaid on the micro-CT scan of the same skull (i) top view, (ii) isometric view (iii) side view from the left, (iv) side view from the right. Scale bar, 1 mm.**(d)** Distribution of measurement error as a function of distance from bregma. **(e)** Cumulative histogram of measurement error for n=6 mice (3 males and n=3 females).

To determine how well the contact sensor could profile the surface of the skull, we profiled the skulls of 6 mice (n = 3 male, n = 3 female C57BL/6J mice) and evaluated the distribution of measurement errors. An x-y point map was used, consisting of 192 points spanning the skull spread across the parietal and frontal bones, covering the whole dorsal cortex. The contact sensing was equipped with either using a needle tipped stylus or a ruby sphere tipped stylus. Five complete scans were performed on each mouse and the average z-coordinate was calculated at each of the measured 192 points. These coordinates were used to construct an average x-y-z point cloud. This point cloud made a conforming fit when superimposed on a 3D surface generated via a micro-computed tomography (CT) scan of the skull of the same mouse (**Fig. 2c**). This was a qualitative indication that the contact sensor could accurately measure the topology of the dorsal skull surface.

In order to quantify the precision of the contact sensor, we calculated, for each point, the measurement error and examined its variation as a function of the distance from bregma (**Fig. 2d**). The maximum measurement error across all measurements was 11 μm (n= 6 mice, 3 males and 3 females). Over 97% of the measurements had an error of less than 6 μm **(Fig. 2e)**. The average measurement error was 1.7 ± 1.4 μm (mean ± standard deviation) for female mice (n=3 mice), and 2.2 ± 1.8 μm for male mice (n=3 mice). There was no significant difference in the mean error between male and female mice (p = 0.62, t-test). Qualitatively, higher error was observed at lateral extremes on the skull surface. The effect of the instantaneous slope of the skull on the measurement error was examined, particularly with the ruby sphere stylus which is more susceptible to forces acting at an angle to the normal axis. There was a slight correlation between the measurement error and the instantaneous slope on the skull surface (r = 0.18, **Supplementary Fig. 5**). If the Craniobot needs to be used on surfaces with steeper curvatures than the experiments’ conducted in this study, it is possible to alleviate this issue by incorporating a tilt axis such that the contact sensor approaches the skull normal to the surface it is profiling. Overall, there was no significant difference between the measurement error of the contact sensor when equipped with the ruby sphere stylus tip versus when equipped with the needle stylus tip (p= 0.1288, t-test). Thus, the contact sensor could be used to precisely map most of the skull surface spanning the dorsal cortex.

### Surface profile guided machining

Once robust and accurate means of profiling the surface of the skull were established, we examined how well this information could be used to guide the CNC mill to perform microsurgical procedures. The dimensions of the stylus used for surface profiling and the dimensions of the cutting tool had to be taken into account when performing this study. A milling path was designed to guide a CNC tool to mill 50 μm deep trenches along the anterior-posterior axis at various distances spaced 500 µm from the midline suture (**Fig. 3a**). Surface profiles were generated by equipping the contact sensor with either a needle tip stylus or a ruby-tip. A 200 µm square end mill and a 300 µm ball end mill resembling commonly used dental burs were used for the cutting operation (**Fig. 3b, Table 1**). The combination of the needle tip stylus and the 300 μm ball end mill resulted in an average trench depth of 44.2 ± 5 μm (n = 3 mice, 78 measured points). The average trench depth was 96.4 ± 19 μm for the combination of the needle tip stylus and 200 μm square end mill (n = 1 mouse, 27 measured points). Using the ruby sphere for surface profiling in combination with the 300 μm ball end mill resulted in an average trench depth of 64.0 ± 5 μm (n= 2 mice, 86 measured points). The combination of ruby sphere tip with the 200 μm square end mill resulted in average trench depth of 57.3 ± 6 μm (n = 3 mice, 77 measured points) (**Fig. 3c**). The trench depth measurements were taken at randomly chosen locations, independent of the locations of the pilot points.

**Figure 3.**
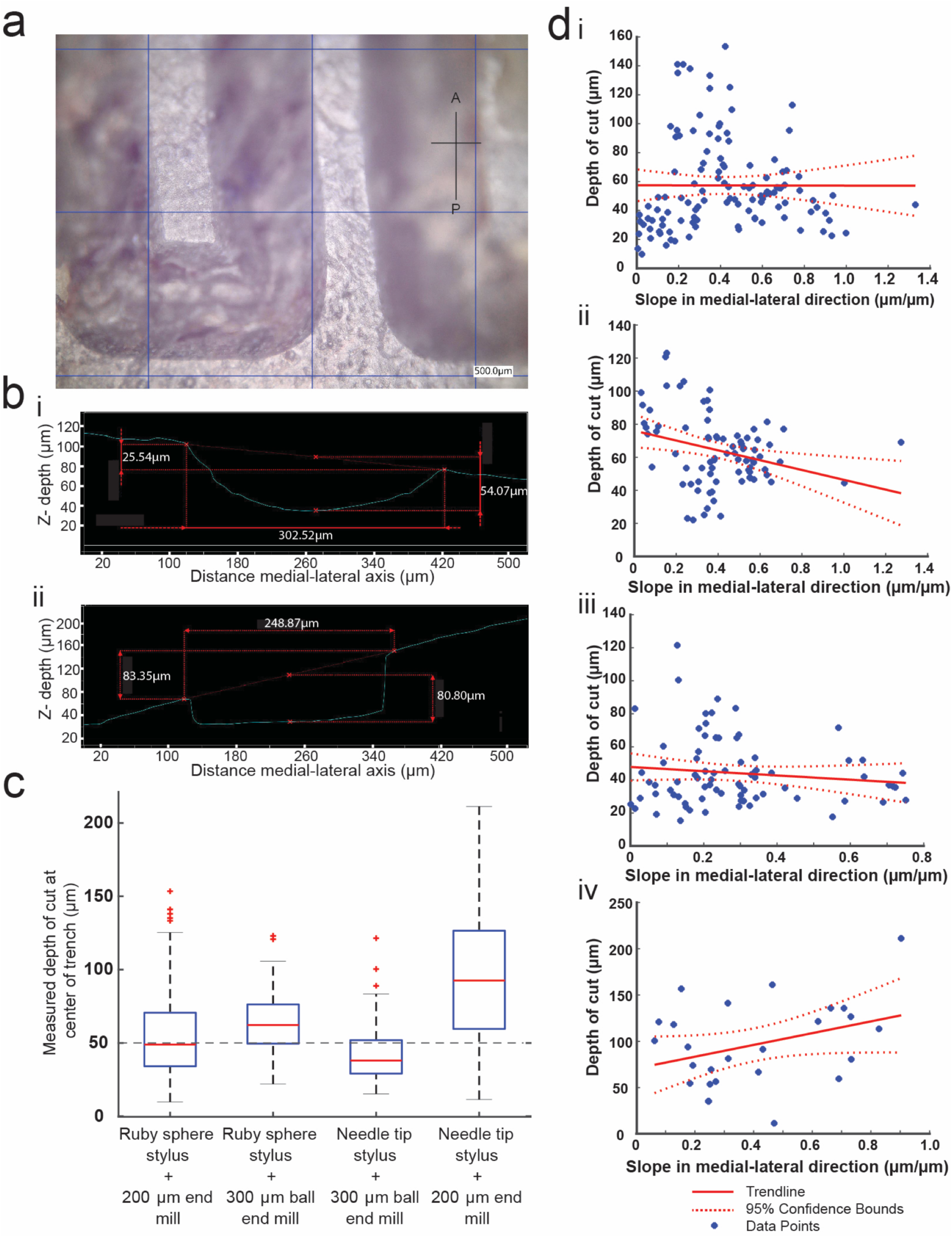
Characterizing performance of surface profile guided CNC machining of the skull: **(a)** Photomicrograph of the 50 μm deep test milling path on the right hemisphere ∼4 mm posterior to bregma and ∼1.5 mm lateral to midline, drilled using the 300 μm ball end mill. Grid size, 500μm. **(b)** Cross sectional profile of the trenches drilled with the 300 μm ball end mill (i) and the 200 μm square end mill (ii) constructed from the photomicrographs **(c)** Boxplot of the measured depth of the trenches for each of the four possible combinations of surface profiling and cutting tools. The dashed line represents the target depth of cut, in contrast with the median value for each combination shown in red. **(d)** Linear regression analysis performed to investigate the effect of the skull morphology on the performance of the Craniobot. Plots for different combinations of profiling and cutting tools are shown for (i) surface profiling with ruby sphere stylus followed by milling with 200 μm square end mill, (ii) surface profiling with ruby sphere stylus followed by milling with 300 μm ball end mill, (iii) surface profiling with needle tip stylus followed by milling with 300 μm ball end mill and (iv) surface profiling with needle tip stylus followed by milling with 200 μm square end mill.

**Table 1.** Summary of trench milling depth measurements

| Surface profiler used | Cutting tool used | Average trench depth ($\mu\text{m}$) | # of mice | # of measured points |
| --- | --- | --- | --- | --- |
| Needle tip stylus | 300 $\mu\text{m}$ ball end mill | $44.2 \pm 5$ | 3 | 78 |
| Needle tip stylus | 200 $\mu\text{m}$ square end mill | $96.4 \pm 19$ | 1 | 27 |
| Ruby sphere tip | 300 $\mu\text{m}$ ball end mill | $64.0 \pm 5$ | 2 | 86 |
| Ruby sphere tip | 200 $\mu\text{m}$ square end mill | $57.3 \pm 6$ | 3 | 77 |

The cutting path between the pilot points was linearly interpolated. Much of the skull surface above the dorsal cortex is convex in the anterior posterior direction, thus, such a linear interpolation could lead to trench depths larger than 50 µm. The average cutting depth across all combinations of surface profiling styluses and cutting tools was 59.3 ± 31.7 µm. Three of the four combinations resulted in trench depths close to the desired value of 50 µm, with the needle tip stylus and the 300 μm ball end mill combination performed closest to target. Given that in our preliminary investigation (**Supplementary Note 1**), the milling operation was needed to be stopped on an average 56 μm (**Supplementary Fig. 3**) before reaching the lower surface of the skull for successful bone removal, we concluded that the precision of the Craniobot was sufficient to safely perform a wide range of microsurgical procedures.

Further, we tested whether the instantaneous slope of the skull surface in the medial-lateral direction affected the depth of trench drilling in each of the four conditions (**Fig. 3d)**. Only the combination of ruby sphere probe and the 300 μm ball end mill displayed a significant relationship between the trench depth and the slope in the medial-lateral direction (p = 0.008, linear regression). In the other cases, there was no statistically significant evidence that the slope of the skull had an effect on the depth of cut. Based upon these results, we concluded that the Craniobot is not only capable of precisely mapping the skull surface, but it is also able to perform micrometer resolution machining operations across a wide swathe of the dorsal skull on stereotaxically-fixed mice.

### CNC microsurgical procedures performed using the Craniobot

To demonstrate the precise surgical capabilities of the Craniobot, we performed 3 mm wide circular craniotomies followed by implantation of glass coverslips to expose the primary somatosensory cortical area for chronic imaging (**Fig. 4a**). The initial milling was started at a depth of 50 µm, followed by an inspection of the bone to examine if it was fragile enough to be excised. If not, additional milling operations were performed, with each subsequent milling run having the depth incremented at 10-15 µm. The final drilling depth was 71 ± 13 µm (6 craniotomies in n=4 C57BL/6J mice, with two bilateral craniotomies performed in 1 mouse, one craniotomy in n=1 Thy1-GCaMP6f). For all surface profiling operations, the Craniobot spent 6.99 ± 0.05 seconds for each point in a surface profiling map (n=3 measurements), and each milling operation took 1.2 ±0.1 seconds per point in the milling path (n=7 measurements). Thus, the cutting operation was typically completed within 2-3 milling iterations and took no longer than 4 minutes. The bone could be successfully excised in all cases without causing damage to the underlying tissue. Thus, this is a relatively quick and high-throughput procedure as compared to manual craniotomies. The chronically implanted windows allowed clear optical access for imaging across days and could also be programmed to drill pilot holes for implantation of bone anchor screws for securing implants (**Fig. 4b**).

**Figure 4.**
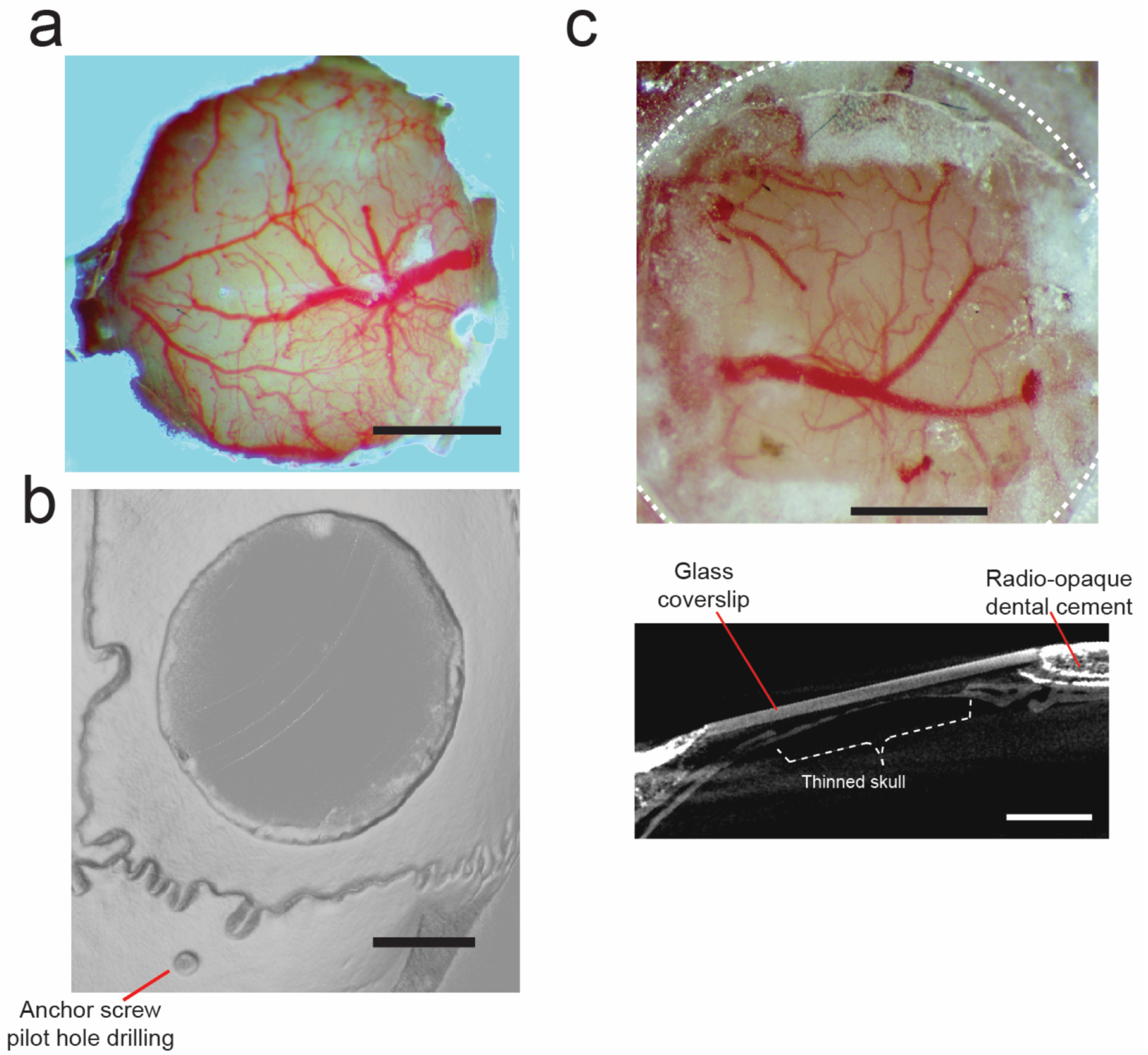
The Craniobot enables us to perform a variety of precise cranial microsurgical procedures in mice. **(a)** A photograph of chronically implanted circular cover-glass in a male C57BL/6J mouse taken 6 days after surgery. Scale bar, 1mm. **(b)**A micro-CT scan of an adult male mouse skull after an automated circular craniotomy was performed and an anchor screw pilot hole was drilled. Scale bar, 1mm. **(c)** Top: A photograph taken after skull thinning operation using the Craniobot, reinforced with clear dental cement and glass coverslip, emphasized by a dashed white line, from an adult male C57BL/6J mouse, 36 weeks old. Surface milling was performed to a depth of 70 μm in a 2 x 2 mm square. Bottom: A micro-CT scan of the same mouse. A cross-sectional view of the thinned portion of the skull implanted with a glass coverslip demonstrating intact skull after thinning. Scale bar, 1mm.

Optical access to the brain can also be gained through the intact skull by thinning and polishing followed by reinforcement with transparent dental acrylic cement (Drew et al., 2010; Shih et al., 2012). The Craniobot was programmed to collect skull surface coordinates from 169 pilot points in a 2 mm x 2mm section to guide surface milling. **Figure 4c** shows a photograph of a mouse skull thinned to depth of 70 μm. Reinforcing the skull with clear dental cement, followed by capping with a glass coverslip allows chronic optical access to the cortex. Thus, the Craniobot can perform intricate microsurgical procedures with precision and repeatability. For simple craniotomies, it provides orders of magnitude increase in speed. While the surface scanning for skull thinning took longer, several parameters such as increasing speed of the contact sensor when approaching the skull, reducing the density total number of pilot points, decreasing the retraction distance after profiling a point and using a milling tool with larger contact area can be explored to optimize this process.

## DISCUSSION

The Craniobot is a generalized, robotic microsurgery platform, which combines automated surface profiling with CNC milling to perform a variety of cranial microsurgical procedures. The Craniobot provides access to the brain in a fast and systematic fashion when compared to current methodologies. We have demonstrated the Craniobot’s ability to perform craniotomies for implantation of chronic cranial windows, and skull thinning for optical imaging through intact skull, and to drill pilot holes for bone anchor screw implantation.

These cranial microsurgery procedures can be performed with micrometer precision in an automated and digitized fashion across the skull independent of the 3D morphology of the skull, without damaging the underlying tissue. Automated microsurgical procedures can be completed within a few minutes resulting in an order of magnitude increase in throughput even when compared to highly experienced surgeons. Thus, the Craniobot is a high-throughput microsurgery platform which lowers entry barriers, reduces training time and cost of performing intricate microsurgical procedures, thereby democratizing the use of advanced neurotechnologies.

In this study, the utility of the Craniobot for performing microsurgical procedures on the skull above the whole dorsal cortex was demonstrated. The same methodology could be extended beyond the parietal and frontal bones above the dorsal cortex for automated microsurgery of the occipital bones above the cerebellum. In the future, we could explore extending the Craniobot to perform bone tissue removal procedures on organs that have much more complex topologies such as the spinal cord. Given the precision of the Craniobot, it could also be extended to smaller animals with delicate skulls such as song birds or juvenile mice.

The Craniobot is fully open-source and uses a high-level machine language, G-code, to automate microsurgical procedures. Given that G-code programming is ubiquitous in manufacturing and precision machining, the programming environment is highly developed, and the Craniobot has access to massive libraries of automation scripts and programs. In the future, more complex microsurgical procedures can be explored by harnessing these resources.

The Craniobot hardware is inexpensive, and the entire setup is built using off-the-shelf components for a fraction of the cost of commercially available stereotaxes. Modifications of the hardware, for instance integrating a tilt axis for machining highly curved surfaces can be easily accomplished within the same framework. In the future, additional functionalities such as bone flap removal, fluidics for irrigating tissue during surgery, control of virus injections, insertion and implantation of devices can all be integrated with relatively simple modifications to the hardware.

Currently, the Craniobot is an open loop system, and its micrometer scale precision is sufficient to perform microsurgical procedures. Closing the operation loop using real-time depth sensing would increase the viability of the Craniobot. Real-time depth sensing, such as optical coherence tomography would enable these automated procedures to be performed in cases where skull thickness cannot be reliably estimated beforehand.

The CNC microsurgery principles can potentially be adapted for larger animal models such as non-human primates. In such settings, access to structural imaging data such as computed tomography (CT) and magnetic resonance imaging (MRI) scans, can be used for depth co-registration with the surface profiling for safe milling procedure. Additionally, force-feedback chucks can be used to automatically disengage the cutting tool in the rare instances when the cutting tool penetrates the bone completely. These advances could allow extending the Craniobot to applications where safety and reliability are critical and could then be translated to clinical use.

Currently, optical imaging is predominantly confined to mice, because of highly developed genetic strategies for expressing optical reporters and optogenetic molecules (Dana et al., 2014; Madisen et al., 2012, 2015; Ting & Feng, 2013; Zeng & Madisen, 2012). With the advent of new genetic tools such as CRISPR, it will be possible soon to easily generate genetically modified rats and nonhuman primates that provide capabilities to study complex cognitive behaviors that are not possible in mice (Chapman et al., 2015; Niu et al., 2014). Generalized strategies to extend cranial microsurgical procedures optimized in mice to these species will be very useful. Automation tools such as the Craniobot can facilitate faster derivation of such procedures.

Robots such as the Craniobot reduce experimental variability and enhance throughput. This could bolster emerging industrial scale neuroscience initiatives such as the Blue Brain project (Markram, 2006), connectomics (Bohland et al., 2009), and Allen Brain Atlases (Oh et al., 2014). By automating cranial microsurgeries, the Craniobot could streamline these neuroscience pipelines and allow neuroscientists to systematically generate vast datasets with ease.

## METHODS

### The Craniobot Construction and Design

#### i. The mill framework

A 3-axis hobby machining mill (Lukcase LC8110) was used as the base machine for the Craniobot because it is widely sold by a variety of manufacturers online, but is generically called a 2020B CNC mill, which refers to the size of the x-y bed of the machine (20cm x 20cm). The frame of the mill was modified to include a home-built custom stereotax in place of the stock mill bed (**Supplementary Fig. 4**). The mill framework is made from machined plastic, and the moving axes are guided along stainless-steel shafts and linear bearings pressed into the framework components. 3D positioning is accomplished by three NEMA 17 stepper motors and M8 stainless steel lead-screw assemblies, all part of the machining mill. The lead-screws couple to each moving axis by anti-backlash nuts, which reduce slop and improve the overall precision of the machine.

#### ii. Custom stereotax

The custom stereotax was constructed by first machining a base out of a 0.75-inch thick aluminum block using a water jet cutter with added abrasives. Holes were water jet cut, drilled, and then tapped into the aluminum base to fit the stereotax pieces and mount it on the mill base. The base was finished with a wire wheel brush to smoothen out the surface. The stereotax parts were made via a combination of 3D printed parts, off-the-shelf components, and machined parts for added stability (**Supplementary Fig. 4**).

#### iii. Control Electronics

We replaced the factory electronics included in the hobby mill with an open-source universal serial bus (USB) stepper motor controller (TinyG v8, Synthetos) that is widely available and has an active online development community. It combines a microcontroller (Atmel ATxmega192) and four stepper motor drivers (TI DRV8811) onto a single printed circuit board and USB interface. An additional microcontroller (Arduino Due, Atmel ATSAM3X8E) was used to connect the contact sensor to the stepper motor controller.

#### iv. Software

Communication to and from the machine was accomplished over USB, and formatted using standardized CNC G-code language packaged inside JavaScript Object Notation (JSON) text strings. The entire robot is operated through a software suite created in Python 3.6 (Python Software Foundation), which generates, sends and receives commands, and communications with the Craniobot. First, the software generates the x-y coordinates of acraniotomy based upon user inputs (i.e. a 3 mm diameter circular craniotomy offset 1.5 mm from bregma, or a whole-dorsal skull craniotomy with a custom shape) and formats these in G-Code-based JSON strings. The Craniobot then uses the customized contact sensor to scan for the z coordinate at each point.

#### v. Contact sensor

A custom contact sensor modified from an industrial digitizing surface probe (Tormach, SPU-40) is monitored via analog input from a microcontroller (Arduino Due). The contact sensor electronics consists of three normally closed switches connected in series. Each switch consists of two spherical stainless-steel contacts electrically bridged by a brass cylindrical arm. The three arms are pressed into the probe tip assembly, and this assembly nominally rests on the three sets of spherical ball contacts creating a normally closed switch circuit. When the probe tip contacts the mouse skull, one of the arms lifts off the spherical contacts and open the circuit. The internal customized spring presses the tip assembly onto the contacts and improves contact resistance. Additionally, we 3D printed a new housing that allowed us to adjust actuation force given by a customized flat spring with #0-80 screws. The resulting design is illustrated in **Figure 2a**.

#### Surgical procedure

All animal experiments were conducted in accordance with protocols approved by the University of Minnesota’s Institutional Animal Care and Use Committee (IACUC). Characterization and demonstration experiments were conducted in acute anesthetized conditions. Mice (C57BL/6J, 8-14 weeks old, Jackson Laboratories) were administered buprenorphine (Par Pharmaceutical, Chestnut Ridge, NY) at 1 mg/kg body weight, and meloxicam (Dechra Veterinary Products, Overland Park, KS) 1-2 mg/kg body weight at the time of the experiment. Mice were anesthetized with 1-5% isoflurane (Piramal Critical Care Inc., Bethlehem, PA), the fur was shaved from the scalp, and the mouse was placed on the custom stereotaxic apparatus. Standard aseptic techniques were used to sterilize the scalp area before the incision. The skin and fascia were then carefully removed around the targeted cutting path. All procedures were performed with aseptic techniques. For characterization experiments, mice were euthanized and perfused immediately upon the completion of the procedure. This was followed by a variety of experiments that were used to characterize the performance of the contact sensor and the ability of the CNC mill to use the surface profiling information in microsurgical procedures, and demonstration experiments.

For microsurgical procedures involving chronic implantations, slow-releasing buprenorphine (Buprenorphine SR LAB, Zoopharm, Windsor, CO) at 1 mg/kg body weight was administered along with meloxicam two hours before the experiment. A 3.3 mm diameter circular craniotomy was performed using the Craniobot, and covered with a 4 mm diameter glass coverslip (Deckgläser, Marienfeld-Superior Inc.) using cyanoacrylate glue (Vetbond, 3M, Saint Paul, MN). Dental cement (S380, C&B Metabond, Parkell Inc.) was then applied to secure the coverslip to the skull surface. We performed chronic implantations of 4 mm glass coverslips in 4 mice using this procedure. After the implantation, the mice were recovered on a heating pad and monitored until they could move autonomously. Meloxicam was administered for 3 days after the experiment along with general monitoring for signs of pain.

### The Craniobot Operation

The Craniobot functions via automating two major processes: CNC profiling the skull surface and using that information to perform CNC milling. Both procedures are described in the following.

#### i. Surface profiling procedure

A flowchart illustrating this procedure is shown in **Supplementary Figure 6**. To profile the skull surface, the needle tip (II M2 030 03 015, ITP Styli LLC, St Louis, MO) or ruby sphere (TH M2 003 03 010, ITP Styli LLC, St Louis, MO) stylus were sterilized with 70% ethanol, washed with 0.9% sterile saline, and attached to the end of the custom contact sensor. The contact sensor was then mounted on the Craniobot’s spindle and was positioned 2-5 mm above the bregma by sending jog commands to the Python software suite. The contact sensor was used to measure the pilot point coordinates in the milling path, which were typically spaced with a maximum distance of 0.5 mm. The Craniobot moved the contact sensor down at a speed of 5 mm/min while constantly sensing the switch signal attached to the sensor. Once a contact was detected, the bregma coordinates were registered as the origin of a Cartesian coordinate system. Once instructed to begin surface profiling along the cutting path, the Craniobot systematically profiled the z coordinate at each pilot point in the desired cutting path.

#### ii. Milling procedure

A flowchart illustrating the milling process is shown in **Supplementary Figure 7**. First, the contact probe was replaced by a cutting tool. We used a 200 μm diameter square end mill (13908, Harvey Tool Inc.), and a 300 μm ball end mill (24512, Harvey Tool Inc.) as cutting tools. The tip of the end mill was carefully brought in contact with the skull surface at bregma, while observing under a stereomicroscope. Then, the experimenter registered bregma as the origin of a new coordinate system used for the milling procedure. The Craniobot then guided the cutting tool through the 3D milling path generated from surface profiling. Saline was used to remove tissue debris during the milling process. At the end of the milling procedure, the cutting tool was raised up 2 mm above the skull surface and stopped. At this point, the experimenter could inspect the engraved/excised skull and decide to stop the milling operation or start another iteration by increasing the milling depth until the milled skull flap could be fractured and excised.

### Perfusion

Following acute surgical procedures, mice were either euthanized or transcardially perfused to preserve the tissues for future analysis. Perfusions were performed by first ensuring the animal was deeply anesthetized with 5% isoflurane. Phosphate-buffered saline (PBS) (CAT# P5493-1L, Sigma Aldrich) was used to flush the circulatory system at a volume of 1 mL/g body weight, and then 4% paraformaldehyde (PFA, CAT# P6148-500G, Sigma Aldrich) was used to fix the tissues. Immediately after the procedure, the skull of the animal was collected and preserved in a solution of 4% PFA.

### Micro-CT scanning

We used micro computed tomography (micro-CT) to analyze and reconstruct the 3D images of the skull after the microsurgical procedures characterization. Perfused mouse skulls were cast in acrylic (Dentsply Orthodontic Resin and Caulk, York, PA, USA) on a Teflon pedestal, and imaged using a Micro-CT machine (XT H 225, Nikon Metrology Inc., Brighton, MI, USA). X-rays were generated with parameters of 105 kV, 85 µA. Mouse skulls were scanned using 720 projections at a half degree pitch while recording 4 frames per projection. Samples were then returned to a solution of 4% PFA after scanning. The micro-CT software reconstructed the x-ray images into a volume graphics (.vgi) file. VG Studio MAX 3.0 (Volume Graphics GmbH, Heidelberg Germany) was used to de-noise unwanted signal caused with a bandpass filter and visualize the 3D structure of the scanned skulls. The 3D structure of the scanned skulls was then transformed into a cloud of 3D coordinates and then exported as a stereolithography (.stl) file.

### Characterizing CNC machining precision

To investigate how the Craniobot’s surface profiling affects the depth of trench milling at different locations on the skull surface, we designed a stereotypical test path. The maximum spacing between pilot points was 0.5 mm, and the trenches were separated by 0.5 mm. Surface profiling was carried out using either the needle-tip or the ruby sphere tip stylus, and milling operation was performed using either the 200 μm square end mill or a 300 μm ball end mill (four conditions). After milling, the mice were immediately perfused, and the skull surface was gently cleaned using sterile saline to remove bone debris from the trenches, after which the skulls were preserved in 4% PFA.

A Digital microscope (Keyence VHX-5000E, at 1500X, 2.1 μm resolution) was used to acquire z-stacks and reconstruct a 3D image of the skull surface at multiple areas. We used the 3D image to calculate the skull slope in the medial-lateral (dz/dx). Measurements were performed every 500 μm, except when residuals such as bone fragments were present.

## ACKNOWLEDGEMENTS

SBK acknowledges funds from the Mechanical Engineering department, College of Science and Engineering, MnDRIVE RSAM initiative of the University of Minnesota, McGovern Institute Neurotechnology (MINT) fund, National Institutes of Health (NIH) 1R21NS103098-01 and 3R21 NS103098-01S1. LG was supported by the University of Minnesota Informatics Institute (UMII) Graduate Research fellowship. GS was supported by the NSF IGERT Neural Engineering traineeship. We would like to thank Bonita Van Heel at Minnesota Dental Research Center for Biomaterials and Biomechanics for help with micro-CT scanning experiments. We would also like to acknowledge the Minnesota Nano Center where we performed the metrology characterizations.

## CONTRIBUTIONS

MR, LG, GJ, ML, JH, GS, DSS and SBK designed, built and tested the automated CNC robots for microsurgical procedures (the auto-driller prototype and the Craniobot). MR, LG, DSS and SBK wrote the manuscript.

## COMPETING FINANCIAL INTERESTS

None.

